# Establishment of a novel amyotrophic lateral sclerosis patient-derived blood-brain barrier model: Investigating barrier dysfunction and immune cell interaction

**DOI:** 10.1101/2024.01.15.575002

**Authors:** Kinya Matsuo, Kazuhiro Nagata, Ryusei Umeda, Takaya Shiota, Satoru Morimoto, Naoki Suzuki, Masashi Aoki, Hideyuki Okano, Masayuki Nakamori, Hideaki Nishihara

## Abstract

Amyotrophic lateral sclerosis (ALS) is a major neurodegenerative disease for which there is currently no effective treatment. The main feature of the disease is the loss of motor neurons in the central nervous system (CNS), but other CNS components and the immune system also appear to be involved in the pathogenesis of the disease. The blood-brain barrier (BBB) is formed by specialized brain microvascular endothelial cells (BMECs) and plays a crucial role in maintaining homeostasis within the central nervous system (CNS). Autopsy studies have reported BBB abnormalities in ALS patients, and studies in animal models suggest that BBB breakdown may precede neurodegeneration. However, the difficulty of obtaining patient samples has hampered further studies. We used the method of differentiating BMEC-like cells from human induced pluripotent stem cells (hiPSCs), which have robust barrier functions and express adhesion molecules necessary for immune cell adhesion, to investigate BBB functions in ALS patients. We also established a method using automated analysis techniques to analyze immune cell adhesion to the BBB in multiple samples simultaneously and with minimal bias. BMEC-like cells derived from ALS patients carrying *TARDBP* ^N345K/WT^ mutations exhibited increased permeability to small molecules due to loss of tight junction, as well as increased expression of cell surface adhesion molecules, leading to enhanced immune cell adhesion. BMEC-like cells derived from hiPSCs with other types of *TARDBP* gene mutations (*TARDBP* ^K263E/K263E^ and *TARDBP* ^G295S/G295S^) introduced by genome editing technology did not show such BBB dysfunction compared to the isogenic control. These results indicate that our model can recapitulate BBB abnormalities reported in ALS patients using patient (*TARDBP* ^N345K/WT^) -derived hiPSCs and that two types of missense mutation, *TARDBP* ^K263E/K263E^ and *TARDBP* ^G295S/G295S^, do not contribute to the BBB dysfunction. This novel ALS patient-derived BBB model is useful for the challenging analysis of the involvement of BBB dysfunction in the pathogenesis and to promote therapeutic drug discovery.

## 1 Introduction

Amyotrophic lateral sclerosis (ALS) is a major and fatal neurodegenerative disease in adults. Primary motor neurons in the cerebral cortex and secondary motor neurons in the anterior horn of the spinal cord are dominantly affected, resulting in progressive limb muscle weakness, dysphagia, and respiratory paralysis(Feldman et al., 2022). The pathomechanism behind motor neuron degeneration remains unknown and there are currently no curative treatments. Although the majority of cases are sporadic, 5-10% of patients have familial forms. There are nearly 30 genes that can cause or modify disease, but the most common pathogenic variants are found in *SOD1, C9ORF72, FUS*, and TAR DNA-binding protein gene (*TARDBP*) (Gregory et al., 2020). *TARDBP* encodes the protein transactive response DNA-binding protein 43(TDP-43), which binds to both DNA and RNA and regulates their expression and splicing. Interestingly, aggregation of TDP-43 in ubiquitin-positive cytoplasmic neuronal inclusions occurs in the brain and spinal cord of ALS patients, including sporadic cases without pathogenic variants in *TARDBP*, and TDP-43 is now recognized as a pathological hallmark of ALS.

Although motor neuron degeneration, a prominent pathological feature of ALS, occurs in the central nervous system (CNS), several studies have reported an association between neurodegeneration and activation of the peripheral immune system and humoral factors in the blood. For example, a cohort study of ALS patients with elevated C-reactive protein (CRP) in peripheral blood showed predominantly impaired motor function and reduced life expectancy (Lunetta et al., 2017). In addition, the population of specific peripheral immune cell subsets in peripheral blood fractions of ALS patients correlated with their functional state and prognosis (Grassano et al., 2023). For instance, elevated peripheral CD8^+^ T cell counts are associated with rapid disease progression (Murdock et al., 2017; Cui et al., 2022). In addition, blood-derived molecules, such as fibrins, and thrombin could be critical elements leading to neuroinflammation and neurodegeneration (Ryu et al., 2018; Shlobin et al., 2021; Finney et al., 2023). Indeed, the presence of serum components and peripheral immune cells within the spinal cord, motor cortex, and brainstem of autopsy samples from ALS patients have been repeatedly reported(Troost et al., 1989; Troost et al., 1990; Kawamata et al., 1992; Engelhardt et al., 1993; Graves et al., 2004; Henkel et al., 2004; Garbuzova-Davis et al., 2012; Winkler et al., 2013; Jara et al., 2019; Garofalo et al., 2020; Garofalo et al., 2022). These data suggest that humoral factors in serum and/or immune cells appear to be important drivers of disease pathogenesis.

The CNS is a sanctuary separated from the peripheral tissues, and there is a strong”barrier” between the two. The blood-brain barrier (BBB) plays a specific role in the brain microvasculature, isolating CNS from its external environment and ensuring its homeostasis. The BBB consists primarily of brain microvascular endothelial cells (BMECs), pericytes embedded in the vascular basement membrane, and astrocyte endfeet aligning the parenchymal basement membrane. BMECs are central to barrier functions, forming continuous tight junctions that prevent water-soluble molecules from diffusing into the CNS via paracellular pathways. They also possess specific polarized transporters and efflux pumps that either take up vital nutrients or remove harmful substances. Importantly, BMECs regulate immune cell migration into the CNS by limiting endothelial adhesion molecules on their surface(Engelhardt and Ransohoff, 2012). Under certain pathological circumstances, the expression of these molecules, such as intercellular adhesion molecule-1 (ICAM-1) and vascular cell adhesion molecule 1 (VCAM-1), is increased, leading to increased and uncontrolled immune cell entry into the CNS.

Although there is growing evidence linking ALS disease progression to activation of the peripheral immune system, key factors that strictly regulate immune cell infiltration into the CNS, such as BBB, have often been ignored in the discussion. Limited evidence has been reported as follows: autopsy studies of ALS patients have shown that extravasated hemoglobin, and plasma-derived proteins such as fibrin, thrombin, and IgG are increased in the spinal cord (Garbuzova-Davis et al., 2012; Winkler et al., 2013). Expression levels of tight junction proteins such as ZO-1, occludin, and claudin-5 are decreased in the medulla and spinal cord of ALS patients (Garbuzova-Davis et al., 2012). These data suggest that the barrier system is impaired in ALS patients. However, it is not well known when this extravasation of serum protein occurs and whether barrier dysfunction contributes to disease onset or progression. In other words, pathological data from ALS patients did not tell us if barrier impairment is just a consequence of neurodegeneration and neuroinflammation or indeed precedes disease onset and relates to disease progression. Although the pathomechanism of BBB alteration in ALS is still unclear, recent studies focusing on *TARDBP* mutations shedding light on the molecular pathway underlying BBB dysfunction in ALS. Some animal studies have suggested that *TARDBP* mutation and/or alteration of TDP-43 expression lead diffusion barrier dysfunction and aberrant immune cell infiltrations (Sasaki et al., 2015; Zamudio et al., 2020; Xu et al., 2022; Cheemala et al., 2023; Hipke et al., 2023). Notably, RNA transcripts and nuclear proteins analysis showed that the brain capillary endothelial cells from patients with ALS or FTLD expressed reduced levels of TDP-43 and nuclear β-catenin along with diminished expression of canonical Wnt response genes, which is important for BBB development and maintenance(Omar et al., 2023). In addition, knockdown of TDP-43 or aberrantly truncated TDP-43 expression in primary human brain endothelial cells results in reduced tight junction protein expressions (Xu et al., 2022; Omar et al., 2023). These reports strongly indicates that TDP-43 plays a regulatory roles in BBB function and physiological dysfunction of TDP-43 is associated with BBB alterations in ALS patients. Despite these increasing interests in relation between TDP-43 and BBB function, there remains a gap in our understanding, as no functional study has yet established a direct link between TDP-43 abnormality and BBB dysfunction using an ALS patient-derived model.

Studying the BBB in patients presents three major challenges. 1) The opportunity to biopsy neural tissue is rare, limiting the ability to observe tissue from ALS at autopsy when end-stage secondary effects cannot be ruled out. 2) Functional evaluations of the barrier cannot be performed using autopsy samples. 3) Studies in animal models make it difficult to reproduce the full pathophysiology, and there are species differences in the important barrier-related proteins that are associated with a lower translation rate to successful human therapy. For this purpose, a BBB model using human-derived tissues that can be functionally analyzed is required. We have recently established the method called extended endothelial cell culture method (EECM) for differentiating BMEC-like cells from human induced pluripotent stem cells (hiPSCs) and investigated the role of the BBB in pathological conditions(Nishihara et al., 2020; Nishihara et al., 2021; Matsuo et al., 2023). Previous studies and reviews reported that hiPSC models reflect human disease pathophysiology, therapeutic effect, and clinical condition of patients (Okano et al., 2020; Okano and Morimoto, 2022; Morimoto et al., 2023; Okano et al., 2023). Widely used differentiation methods for BMEC-like cells(Lippmann et al., 2012; Qian et al., 2017) show a robust diffusion barrier and expression of several BMEC-specific transporters but have some limitations. First, they also express epithelial genes and proteins (Lippmann et al., 2020; Lu et al., 2021). Second, they lack several cell surface adhesion molecules such as selectins, ICAM-2, and VCAM-1, which is essential to studying immune cell interaction with the BBB (Nishihara et al., 2020). Our hiPSC-derived BBB model does not show epithelial genes and possesses both proper tight junction integrity and robust expression of adhesion molecules essential for immune cell interaction with the BBB, making this model unique and suitable for studying the link between the peripheral immune system and the CNS via the BBB (Nishihara et al., 2020; Gastfriend et al., 2021).

In this study, we presented the successful establishment of BMEC-like cells from iPSCs derived from an ALS patient carrying the *TARDBP* ^N345K^ as an *in vitro* ALS-BBB model. Furthermore, we generated a BBB model from two genome-editing hiPSC clones targeting *TARDBP* ^G295S/G295S^ and *TARDBP* ^K263E/K263E^ from the healthy donor-derived hiPSC and investigated whether the specific *TARDBP* mutations contribute to BBB dysfunction. ALS patient-derived but not *TARDBP* ^G295S/G295S^ and *TARDBP* ^K263E/K263E^ gene-edited, EECM-BMEC-like cells showed impaired diffusion barrier functions and enhanced immune cell adhesion compared to control. This model is useful for studying immune cell-BBB interactions and mechanisms how TDP-43 impact on BBB functions, and can be applied to other genetic mutations in the future.

## 2 Material and method

### Donors and human induced pluripotent stem cells (hiPSCs)

The donors of hiPSCs and the origin of the *TARDBP*-mutated gene-editing hiPSCs were summarized in Figure 1A. A patient-derived hiPSC line was reprogrammed from T lymphocytes derived from a patient with familial ALS (age/sex: 63/M, *TARDBP* p.N345K) (Leventoux et al., 2020). The isogenic hiPSC lines that induced two distinct pathogenic mutations of the TDP-43 protein, TDP-43 TDP-43^K263E/K263E^ and TDP-43 ^G295S/G295S^, were generated as described (Imaizumi et al., 2022). Both of the mutations have been reported to be associated with ALS and frontotemporal lobar degeneration (FTLD) (Floris et al., 2015) and the locus of each mutation is shown in Figure 1B. Two healthy controls (HC) derived hiPSC lines HPS1006 and HPS4290 (Takahashi et al., 2007) (age/sex: 36/M, 36/F, respectively) were provided by the RIKEN BRC through the National Bio-Resource Project of the MEXT/AMED, Japan. TDP-43 ^K263E/K263E^ and TDP-43 ^G295S/G295S^were generated from HPS4290. In this report, HPS1006 and HPS4290 are referred to as HC1 and HC2, respectively. Also, hiPSC lines carrying the *TARDBP* ^K263E/K263E^ and *TARDBP* ^G295S/G295S^ mutations are referred to as ALS/FTLD1 and ALS/FTLD2, respectively.

**Figure 1.**
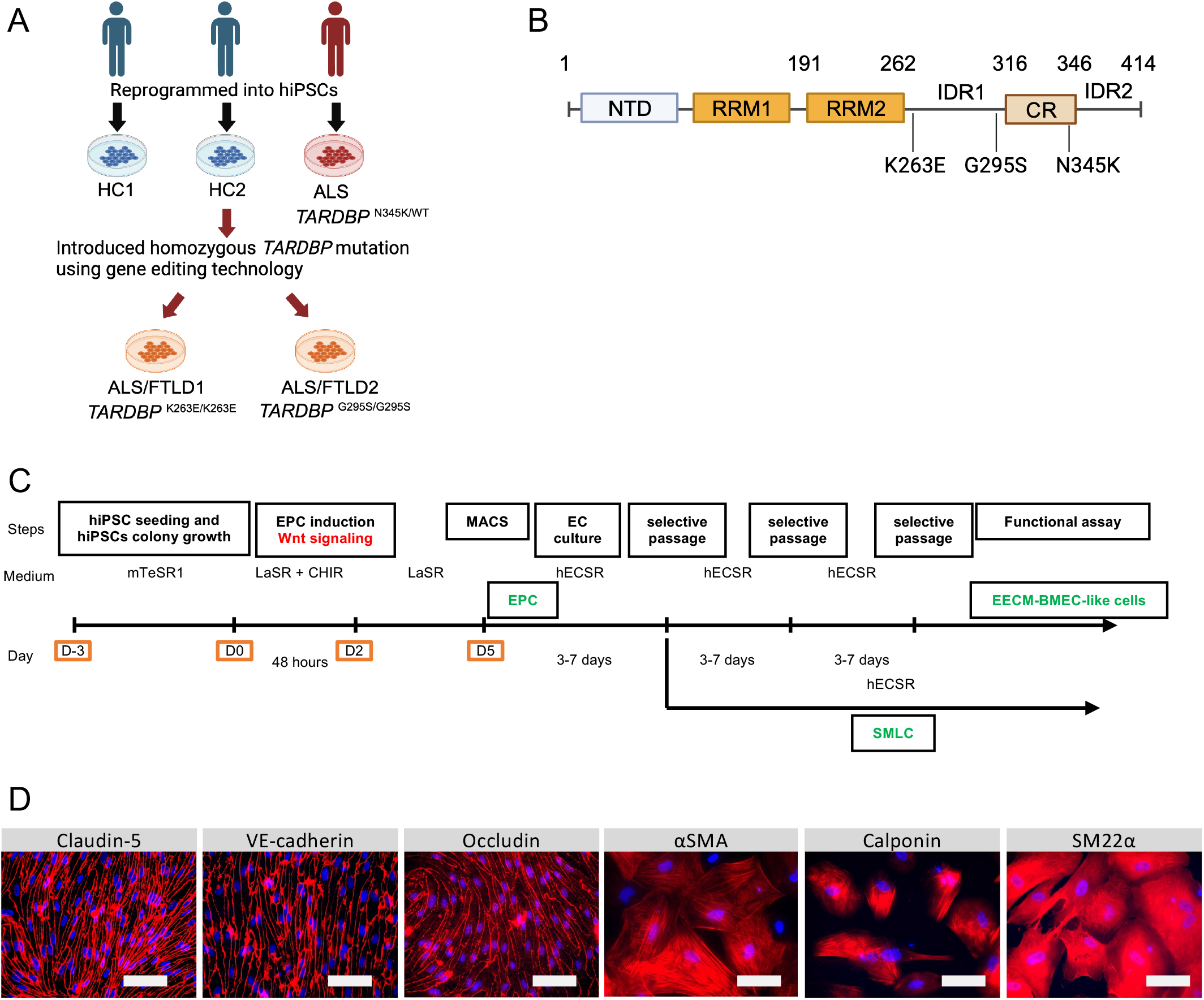
Establishment and characterization of EECM-BMEC-like cells and SMLCs. (A)Overview of the donors of hiPSCs and the origin of *TARDBP* mutated gene editing hiPSC clones are shown. hiPSCs were generated from two distinct healthy donors (HC1 and HC2) and one ALS patient carrying heterozygous *TARDBP* mutation, TDP-43 ^N345K/WT^ (ALS). HC2 is the isogenic control of *TARDBP* ^K263E/K263E^ (ALS/FTLD1) and *TARDBP* ^G295S/G295S^ (ALS/FTLD2). hiPSC: human induced pluripotent stem cells, HC : healthy control, ALS : amyotrophic lateral sclerosis. Figure created with BioRender.com (B) Locus of each *TARDBP* mutations are shown. NTD: N-terminal domain, RRM: RNA recognition motif, IDR: intrinsically disordered regions; CR: conserved region. Figure created with BioRender.com (C) The overview of EECM protocol. EPC : endothelial progenitor cell, MACS : magnetic activated cell sorting, EC : endothelial cell, mTeSR1/LaSR/CHIR/hECSR : specific name of the medium and compound. Recipes are shown in Supplementary table 2. EECM : extended endothelial cell culture method, BMEC : brain microvascular endothelial cell, SMLC : smooth muscle like cell (D) Representative immunostaining of EECM-BMEC-like cells and SMLCs. Cells from HCs were grown in 8-well chamber plate and EECM-BMEC-like cells were stained for junctional protein VE-cadherin, claudin-5 or occludin (red). SMLCs were stained for smooth muscle cell markers αSMA, calponin or SM22α (red). Nuclei were stained using DAPI (blue). Scale bar = 50 μm

### hiPSCs maintenance

hiPSCs were maintained in mTeSR1 medium (STEMCELL Technologies, Vancouver, Canada). When cells became 80% confluency, the hiPSCs were detached using ReLeSR™ (STEMCELL Technologies, Vancouver, Canada) according to manufacture protocol. The cramps of hiPSCs were seeded in another Matrigel-coated six-well plate. Cultural media were changed every day.

### Differentiation of hiPSCs into EECM-BMEC-like cells and smooth muscle-like cells (SMLCs)

The extended endothelial cell culture method (EECM) was employed as an iPSCs-derived human *in vitro* BBB model as previously described (Nishihara et al., 2020; Nishihara et al., 2021; Matsuo et al., 2023) (Figure 1C). CD31^+^ endothelial progenitor cells were generated by stimulating with CHIR99021 which activates Wnt/β-catenin signaling. To obtain a higher purity of CD31^+^ cells, the seeding density of hiPSCs on day -3 was optimized for each hiPSC clone with a range of 24,000/cm^2^ to 53,000/cm^2^(supplementary table 1). After purifying the CD31^+^ cells using magnetic activated cell sorting (MACS), the cells were seeded on a collagen-coated 6-well plate. After 3-5 days of cultivating these cells, a small population of SMLCs is interspersed among the endothelial cells. When these cells are subjected to Accutase™ (Sigma-Aldrich, St. Louis, US) treatment, the endothelial cells detach preferentially due to the differential detachability between endothelial and SMLCs. This difference can be exploited to selectively collect the initially detached endothelial cells. By repeating this process 2-3 times, a pure population of endothelial cells can be obtained and termed EECM-BMEC-like cells. EECM-BMEC-like cells from passages 3 to 6 were used for the following assays. The SMLCs remaining in the 6-well plate are left in the hECSR medium (supplementary table 2), and the conditioned medium is collected every 2-3 days when the medium changes, then filtered through a 0.22 μm filter and stored as previously described (Nishihara et al., 2021; Matsuo et al., 2023).

### Permeability assay

Permeability of the EECM-BMEC-like cell monolayer was measured as previously described (Nishihara et al., 2021). In brief, EECM-BMEC-like cells were seeded onto 0.4μm pore Transwell™ filter inserts (Corning 3401, 12 wells 0.4μm pore size, PC membrane, Corning, US) with a consistent density of 1.12 × 10^5^ cells per 500 μl and cultured for 6 days. On day 6, the monolayers’ permeability was evaluated by determining the sodium fluorescein (NaFl, 376.3 Da, Sigma-Aldrich, St. Louis, US) clearance. A10 μM concentration of NaFl was introduced to the top section of the Transwell™ inserts. Samples with the diffused fluorescent tracer were collected from the lower chamber every 15 minutes for up to 1 hour. The fluorescence intensity was measured using a FlexStation™ 3 multi-well reader (Molecular Devices, San Jose, US), with the excitation set at 485 nm and the emission detected at 530 nm. Each condition underwent the tests in triplicate. After each assay, the filters were stained as described below to confirm that the EECM-BMEC-like cell monolayer did not have holes that could result in incorrect permeability.

### Flow cytometry analysis for cell surface expression of adhesion molecules

Cell surface expression of endothelial adhesion molecules was investigated as previously described (Nishihara et al., 2021). EECM-BMEC-like cells were cultured on 24-well plates with conditioned medium derived from SMLCs of the same clone in the presence or absence of stimulation with 1 ng/ml recombinant human TNF-α (R&D Systems, 210TA) and 20 IU/ml recombinant human IFN-γ (285IF, R&D Systems, Minneapolis, US) for 16 hours at 37°C, 5% CO_2_. Stainings of cell surface adhesion molecules were detected using Attune NxT Flow Cytometer (Thermo Fisher Scientific, Waltham, US). Delta mean fluorescence intensities (ΔMFI: MFI staining – MFI isotype) were calculated by using FlowJo™ Ver10.9 (BD, Franklin Lakes, US).

### Immunofluorescence stainings

EECM-BMEC-like cell monolayers on the filter insert membranes or collagen type 4 coated 8-well chamber plate were analyzed for tight junction proteins as previously described(Nishihara et al., 2021). Briefly, cells were fixed with cold methanol (−20°C) for 20 seconds. Then, cells were blocked and permeabilized with a solution of 5% skimmed milk and 0.1% Triton X-100, followed by incubation with primary and secondary antibodies (indicated in supplementary table 3). The nuclei were stained with 4’,6-diamidino-2-phenylindole (DAPI) at a concentration of 1 μg/ml. After DPBS washes, they were mounted with Mowiol (Sigma-Aldrich, St. Louis, US). Images were captured using a BZ-X810 microscope (KEYENCE, Osaka, Japan).

### Immune cell adhesion assay under static conditions on a 96-well plate

EECM-BMEC-like cells are seeded onto 10 μg/ml collagen type IV-coated 96-well plates (Coning 3596, Coning, US) at a density of 30,000 cells per well. The following day, the medium was replaced with a conditioned medium of SMLCs derived from the same hiPSC clone and incubated for 16 hours in the presence or absence of 0.1 ng/ml TNF-α and 2 IU/ml IFN-γ. On the day of the experiment, fluorescently-labeled PBMCs were prepared and adhesion assays were performed using a previously described method (Nishihara et al., 2021), with slight modifications. Briefly, PBMCs were isolated from healthy donors using BD Vacutainer CPT Mononuclear Cell Preparation Tube (BD, Franklin Lakes, US) as manufacture protocol. Isolated PBMCs were frozen until use. PBMCs from allogeneic healthy donors were thawed on the day of experiments using T-cell wash buffer (supplementary table 2) and were centrifuged (280g, 5min, 20-25°C) to remove the preservation reagent. PBMCs were incubated with the T-cell medium (supplementary table 2) containing 1 nM CMFDA dye (Invitrogen™, Waltham, US) for 30 minutes at 37°C, 5% CO_2_. Unconjugated fluorophores were washed out and labeled PBMCs were resuspended in the T-cell medium (supplementary table 2). Ficoll-Paque PLUS (Sigma-Aldrich, St. Louis, US) was carefully added underneath the cell suspension and the tube was centrifuged (770g, 20 min at 20–25°C, minimum brake and acceleration) and live PBMCs existed in the interphase between the buffer and the Ficoll-Paque were collected and resuspended with the migration assay medium (supplementary table 2) at a concentration of 2.0×10^5^/ml. 100 μL of PBMC suspension (2.0×10^4^/well) was evenly added using electronic multi-dispenser pipettes (Eppendorf, Hamburg, Germany) to each well of a 96-well plate on top of EECM-BMEC-like cells. The plate was placed on a plate shaker with light shading. After 15 minutes, the plate was rotated 90 degrees and incubated for another 15 minutes on the shaker. After sucking up the unattached cells, 200 μL of DPBS (Thermo Fisher Scientific, Waltham, US) was gently added to each well using electronic multi-dispenser pipettes, the washed solution was aspirated. Washing wells was repeated one more time and then fixed cells with 50 μL cold methanol (−20°C) for 1 minute. After washing the wells with 100 μL of DPBS, 100 μL of Mowiol was added to each well. Images of the wells were captured using a BZ-X810 microscope (KEYENCE, Osaka, Japan) at a magnification of 10x. To mitigate the potential for selection bias regarding the photograph’s location, the setup was calibrated to ensure automatic capture of each well’s center. Before analyzing the number of adhered PBMCs, phase contrast images were used to exclude wells in which a part of the EECM-BMEC-like cell monolayer was detached. The number of adherent CMFDA-labeled PBMCs was automatically counted in each captured image using the hybrid cell count software BZ-X analyzer in each visual field (KEYENCE, Osaka, Japan).

### Statistical analysis

GraphPad Prism 7 software (GraphPad Software, La Jolla, US) was used for statistical analyses that included degree of freedom calculations. The displayed data is the mean ± SD. An unpaired t-test was used to determine statistical significance when comparing two groups. The corresponding figure legends provide information on the specific statistical methods applied to each experiment.

### Study approval

The study was approved by the ethics committees of the Medical Faculties of Yamaguchi University (number H2021-133-3) and Keio University School of Medicine Ethics Committee (approval no.20080016).

## 3 Results

### Establishment and characterization of EECM-BMEC-like cells and SMLCs

We first differentiated EECM-BMEC-like cells and SMLCs to establish *in vitro* BBB models from an ALS patient. One hiPSC clone from an ALS patient with *TARDBP* ^N345K^ (ALS) and two healthy controls (HC1, HC2) was used in the study. In addition, by introducing different *TARDBP* mutations into HC2, two distinct *TARDBP* mutant iPSC clones (*TARDBP*^G295S/G295S^ (ALS/FTLD1) and *TARDBP* ^K263E/K263E^ (ALS/FTLD2)) were generated on the same genetic background (Figure 1A). The HC- and ALS-derived iPSCs were differentiated BMEC-like cells and SMLCs using EECM (Figure 1C). Both HC- and ALS-derived hiPSCs can be differentiated into EECM-BMEC-like cells, which resemble primary human BMECs in spindle-shaped morphology (Figure 1D). We further confirmed the continuous adherens junction in the staining for VE-cadherin and the intercellular localization of tight junction proteins such as claudin-5 and occludin (Figure 1D). SMLCs derived from each hiPSC clone also confirmed characteristics of smooth muscle cells by staining such as alpha-smooth muscle actin (αSMA), smooth muscle protein 22-alpha (SM22α), and calponin (Figure 1D). Differentiation of each clone was performed at least 5 times, and its reproducibility was confirmed. These data indicate that both HC- and ALS-derived hiPSCs successfully differentiated into EECM-BMEC-like cells and SMLCs as previously reported (Nishihara et al., 2020) and used further studies for diffusion barrier functions and immune cell interaction with BBB.

### ALS patient-derived EECM-BMEC-like cells, but not *TARDBP* ^K263E/K263E^ and *TARDBP*^G295S/G295S^-carrying cells, showed loss of tight junctions

Since autopsy samples from the brains or spinal cords of ALS patients showed decreased expression levels of tight junction proteins(Garbuzova-Davis et al., 2012), we next examined if ALS patient-derived EECM-BMEC-like cells recapitulate disrupted tight junction. We compared the morphology and the staining of adherence and tight junctions of ALS-derived EECM-BMEC-like cells to HCs-derived cells. As a result, ALS-derived EECM-BMEC-like cells had a larger morphology despite the same seeding density (Figure 2). There was no significant difference in the staining of VE-cadherin, the adherens junction, between ALS-derived EECM-BMEC-like cells and HCs-derived cells. Interestingly, we found that ALS-derived cells showed discontinuation of tight junction proteins claudin-5 and occludin, which was not observed in HCs-derived cells. Next, we asked whether ALS/FTLD1- and ALS/FTLD2-derived cells, which are single *TARDBP* mutant models, recapitulate the disrupted tight junctions. Contrary to expectations, ALS/FTLD1- and ALS/FTLD2-derived EECM-BMEC-like cells showed continuous tight junctions comparable to HC-derived cells (Figure 2). These results indicate that ALS patient-derived EECM-BMEC-like cells mimic the tight junction discontinuation observed in ALS patients’ autopsied CNS samples. Further, the specific *TARDBP* mutations (*TARDBP* ^K263E/K263E^ and *TARDBP* ^G295S/G295S^) do not contribute to disrupted tight junctions.

**Figure 2.**
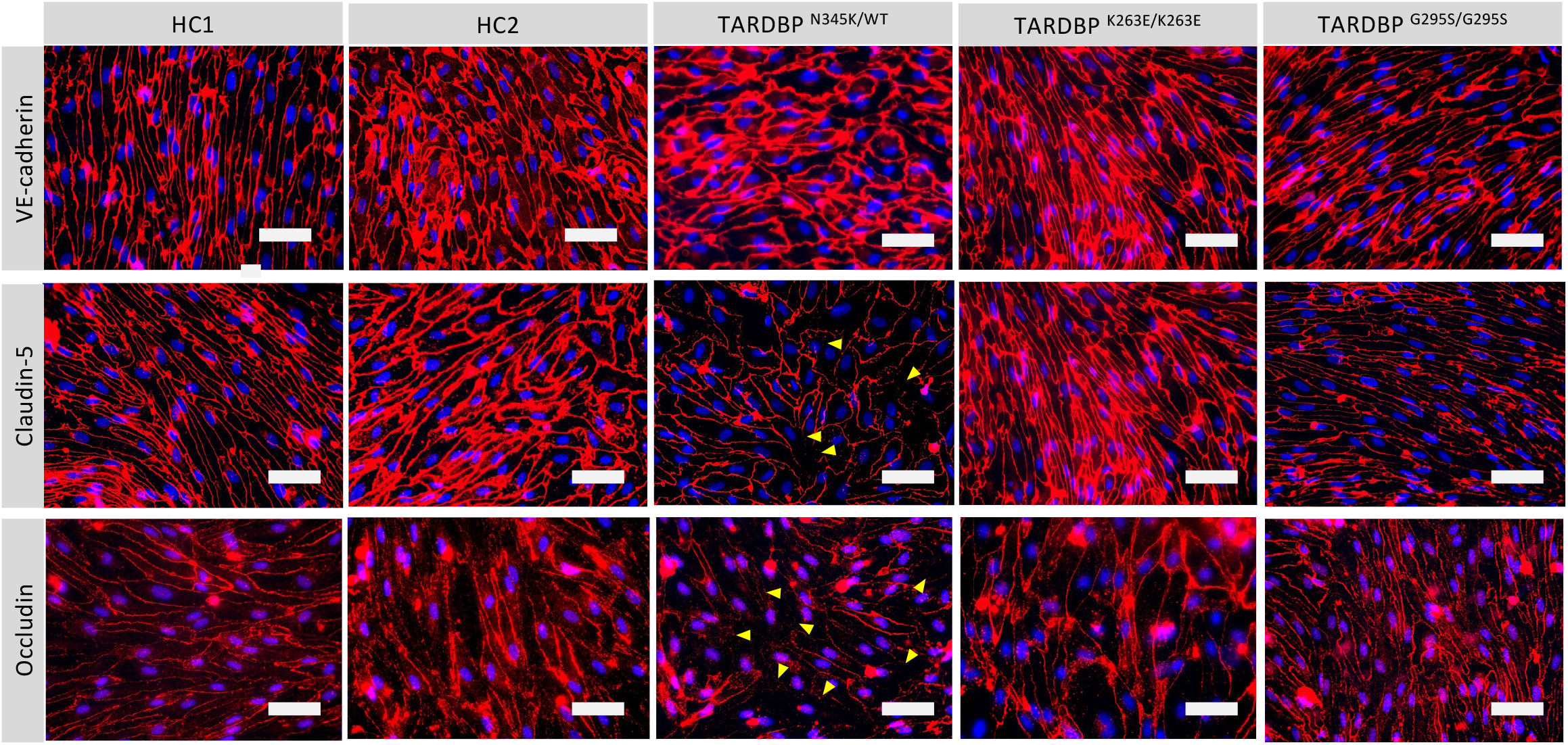
ALS patient-derived EECM-BMEC-like cells exhibit tight junction discontinuation. EECM-BMEC-like cells derived from HC or ALS/FTLD were cultivated onto 0.4um pore size Transwell® filter for six days. Junctions were stained for VE-cadherin, claudin-5 or occludin (red), and nuclei were stained with DAPI (blue). Yellow arrow heads indicate visible discontinuations of the claudin-5 and occludin. Scale bar = 50 μm

### ALS patient-derived EECM-BMEC-like cells, but not *TARDBP* ^K263E/K263E^ and *TARDBP* ^G295S/G295S^-carrying cells, showed impaired diffusion barrier properties

The leakage of blood proteins such as IgG and fibrin from blood vessels has been reported in autopsied brains and spinal cords of ALS patients (Winkler et al., 2013). Still, it was not clear whether this leakage was a result of the end-stage neurodegeneration and postmortem changes or indicated a primary abnormality in the BMECs. To answer this question, we asked if ALS patient-derived *in vitro* BBB model shows increased permeability in the absence of neurodegeneration or inflammation. Comparing permeability for small molecule tracers, as a marker for diffusion barrier properties, ALS-derived EECM-BMEC-like cells showed significantly increased NaFl permeability compared to HCs (Figure 3). Next, we asked whether the specific *TARDBP* mutations (*TARDBP* ^K263E/K263E^ and *TARDBP* ^G295S/G295S^) contribute to impaired diffusion barrier properties. The permeability of ALS/FTLD1- and ALS/FTLD2-derived EECM-BMEC-like cells was not significantly increased compared to HC2-derived cells (Figure 3). These data indicate that EECM-BMEC-like cells derived from an ALS patient (ALS) exhibit impaired junctional integrity and this discontinuation of tight junctions (Figure 2) has functional significance as reflected in the permeability of small molecules (Figure 3). This means that ALS patients show defective diffusion barrier properties in the absence of the factor of neurodegeneration and neuroinflammation. Further, the specific *TARDBP* mutations (*TARDBP* ^K263E/K263E^ and *TARDBP* ^G295S/G295S^) do not contribute to defective diffusion barrier properties.

**Figure 3.**
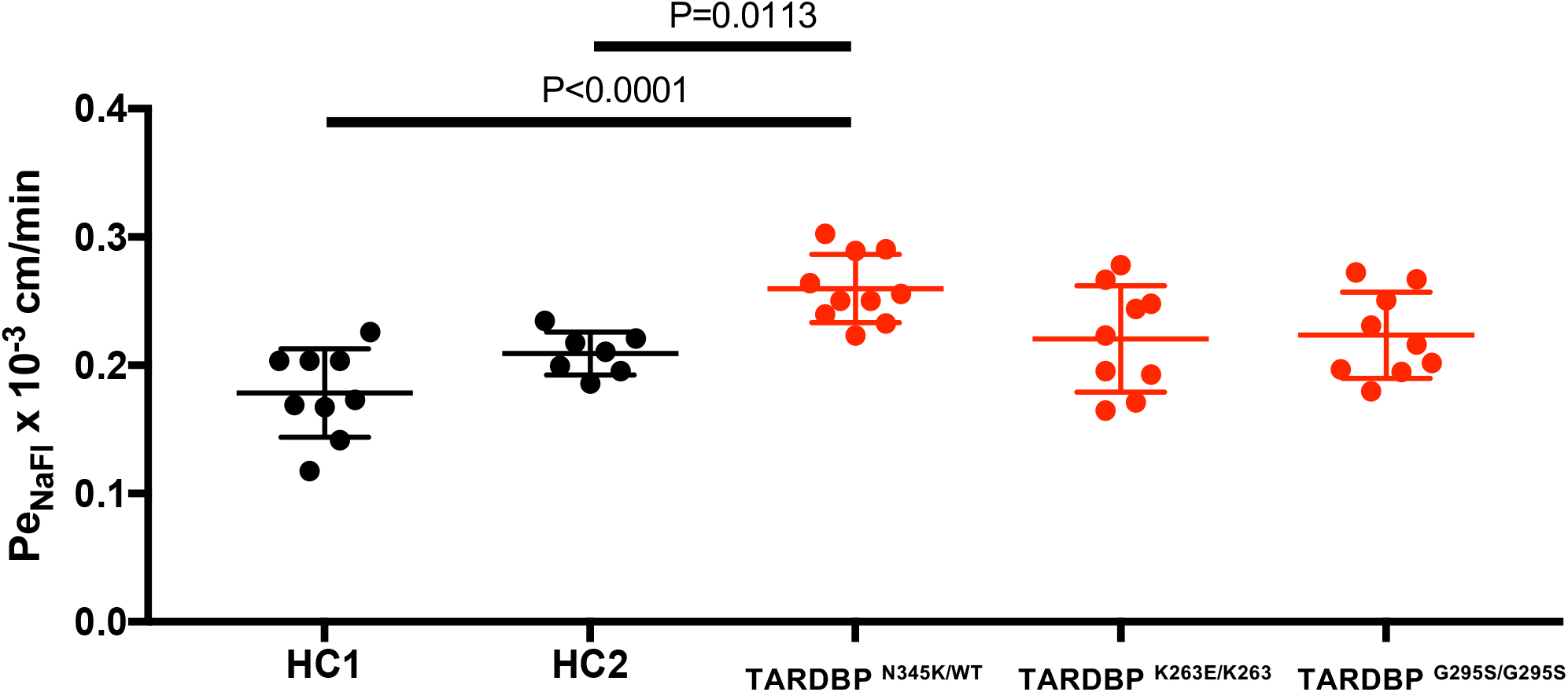
EECM-BMEC-like cells derived from ALS patients exhibited impaired diffusion barrier properties, unlike cells carrying only the *TARDBP* mutation. EECM-BMEC-like cells derived from HC (black) or ALS clones (red) were grown on 0.4 μm pore size Transwell® filters for 6 days to form confluent monolayer in monoculture. Permeability was measured at day 6. The permeability was calculated from the fluorescence intensity of 0.37 kDa sodium fluorescein (NaFl) across the EECM-BMEC-like cell monolayers. Each symbol (HC: black, ALS: red) shows the permeability of independent filters with at least three single differentiations, each performed in at least triplicate. Data are shown as mean ± SD. Statistical analysis: unpaired *t* test. P-values are indicated in the corresponding figures.

### Increased expression of adhesion molecules in ALS patient-derived EECM-BMEC-like cells, but not *TARDBP* ^K263E/K263E^ and *TARDBP* ^G295S/G295S^-carrying cells

ALS-patient autopsy samples show immune cell infiltrations into the CNS (Henkel et al., 2004; Garofalo et al., 2022) as another BBB-disrupting feature. Therefore, we next checked if ALS patient-derived EECM-BMEC-like cells possess important endothelial adhesion molecules ICAM-1 and VCAM-1, which controls immune cell infiltration into the CNS (Marchetti and Engelhardt, 2020) and this model could be used for immune cell-BBB interactions. We examined the expression of cell surface endothelial adhesion molecules in conditions with and without proinflammatory cytokine stimulation, as a representative of physiological and peripheral inflammation-stimulated conditions, respectively. When culturing EECM-BMEC-like cells in an SMLC-conditioned medium, ALS-derived EECM-BMEC-like cells expressed cell surface ICAM-1 under nonstimulated conditions, which is enhanced upon proinflammatory cytokine stimulation (Figure 4A). Cell surface VCAM-1 expression is minimal without cytokine stimulation but is effectively induced upon proinflammatory cytokine stimulation in the presence of SMLC-conditioned medium, as previously described (Nishihara et al., 2020) (Figure 4B). We next compared cell surface adhesion molecule expression in ALS-derived EECM-BMEC-like to HCs-derived cells. We found that ALS-derived EECM-BMEC-like cells expressed increased ICAM-1 under both unstimulated and stimulated conditions and VCAM-1 under stimulated conditions compared to HCs (Figure 4A, B). Next, we asked if ALS/FTLD1- and ALS/FTLD2-derived cells, showed increased expression of endothelial adhesion molecules in both unstimulated and stimulated conditions compared to HC2. Surprisingly, ALS/FTLD1- and ALS/FTLD2-derived cells didn’t show enhanced expression of cell surface ICAM-1 and VCAM-1. These data suggest that ALS patient-derived EECM-BMEC-like cells express enhanced ICAM-1 and VCAM-1, which may contribute to immune cell trafficking into the CNS, but again, the specific *TARDBP* mutations (*TARDBP* ^K263E/K263E^ and *TARDBP* ^G295S/G295S^) do not contribute to the increased cell surface adhesion molecules.

**Figure 4.**
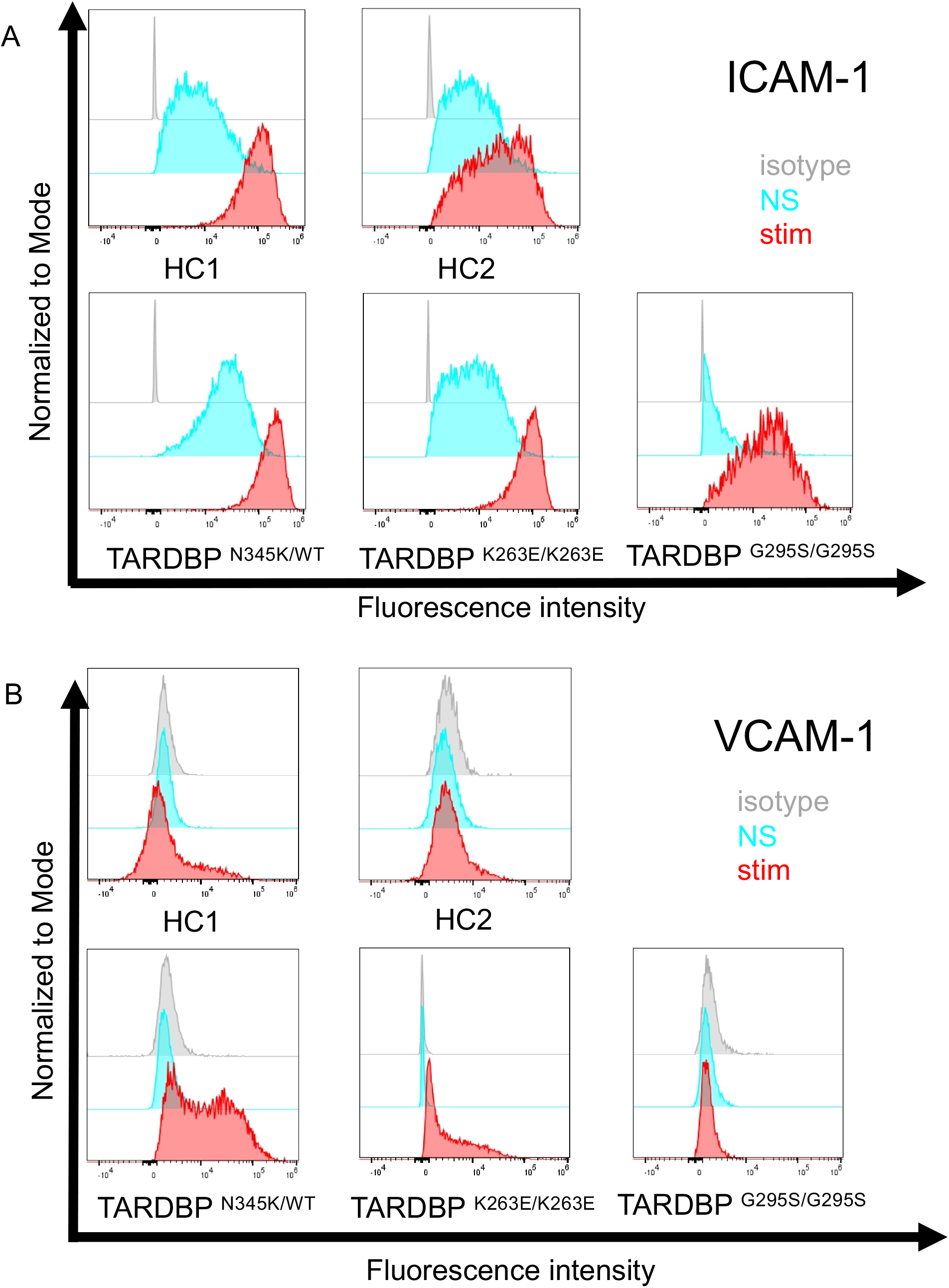
ALS patient-derived EECM-BMEC-like cells showed enhanced endothelial adhesion molecules. EECM-BMEC-like cells are seeded onto type IV collagen-coated well plates and treated with conditioned medium from the same hiPSC clone, with or without proinflammatory cytokine stimulation (1 ng/ml TNF-α + 20IU/ml IFN-γ). Flow cytometry was used to detect cell surface immunostaining of EECM-BMEC-like cells for the adhesion molecules ICAM-1 and VCAM-1. Gray, blue, and red lines represent isotype control, non-stimulated (NS), and 16-hour proinflammatory cytokine-stimulated conditions, respectively. (A) Representative histograms for ICAM-1 are shown for each clone. At least three replicates were analyzed for each clone. (B) Representative histograms for VCAM-1 are shown for each clone. At least three replicates were analyzed for each clone.

### ALS patient-derived EECM-BMEC-like cells enhance PBMC adhesion

Finally, we investigated whether the adhesion molecules on the surface of EECM-BMEC-like cells are functional and enhanced cell surface ICAM-1 and VCAM-1 in ALS-derived EECM-BMEC-like cells are correlated to increased immune cell interaction with BBB. To this end, the immune cell adhesion assay was performed and the number of adherent allogeneic PBMCs on EECM-BMEC-like cells monolayer from each clone was compared. This assay is done based on the previously reported methodology (Nishihara et al., 2021) with modification. EECM-BMEC-like cells from all clones were seeded in a 96-well plate and assayed in the same setup to minimize the variability between the experiments (Figure 5A). To avoid selection bias, the center of all wells was automatically photographed. Although the cells in all wells formed a complete monolayer before the assay, some of the cells were detached after the assay due to the fixation and washing process, therefore, we excluded these wells from the analysis using phase contrast images. The number of adherent PBMCs was then counted in the same setup concerning the fluorescence threshold for image analysis using the hybrid cell count software BZ-X analyzer. The fluorescently labeled PBMCs dispersed well and adhered to the monolayer of EECM-BMEC-like cells in each well (Figure 5B). When we compared the number of adherent PBMCs, ALS-derived EECM-BMEC-like cells showed a significantly increased number of adherent PBMCs under proinflammatory cytokine-stimulated conditions compared to HCs (Figure 5C). In contrast, ALS/FTLD1- and ALS/FTLD2-derived EECM-BMEC-like cells didn’t show such enhanced PBMCs attachment compared to HC2. Thus, the elevated levels of cell surface ICAM-1 and VCAM-1 in EECM-BMEC-like cells derived from ALS patient is functional in attracting more immune cells. Furthermore, this enhanced PBMC adhesion to the ALS-derived BBB model happens independent of changes within the CNS including neurodegeneration.

**Figure 5.**
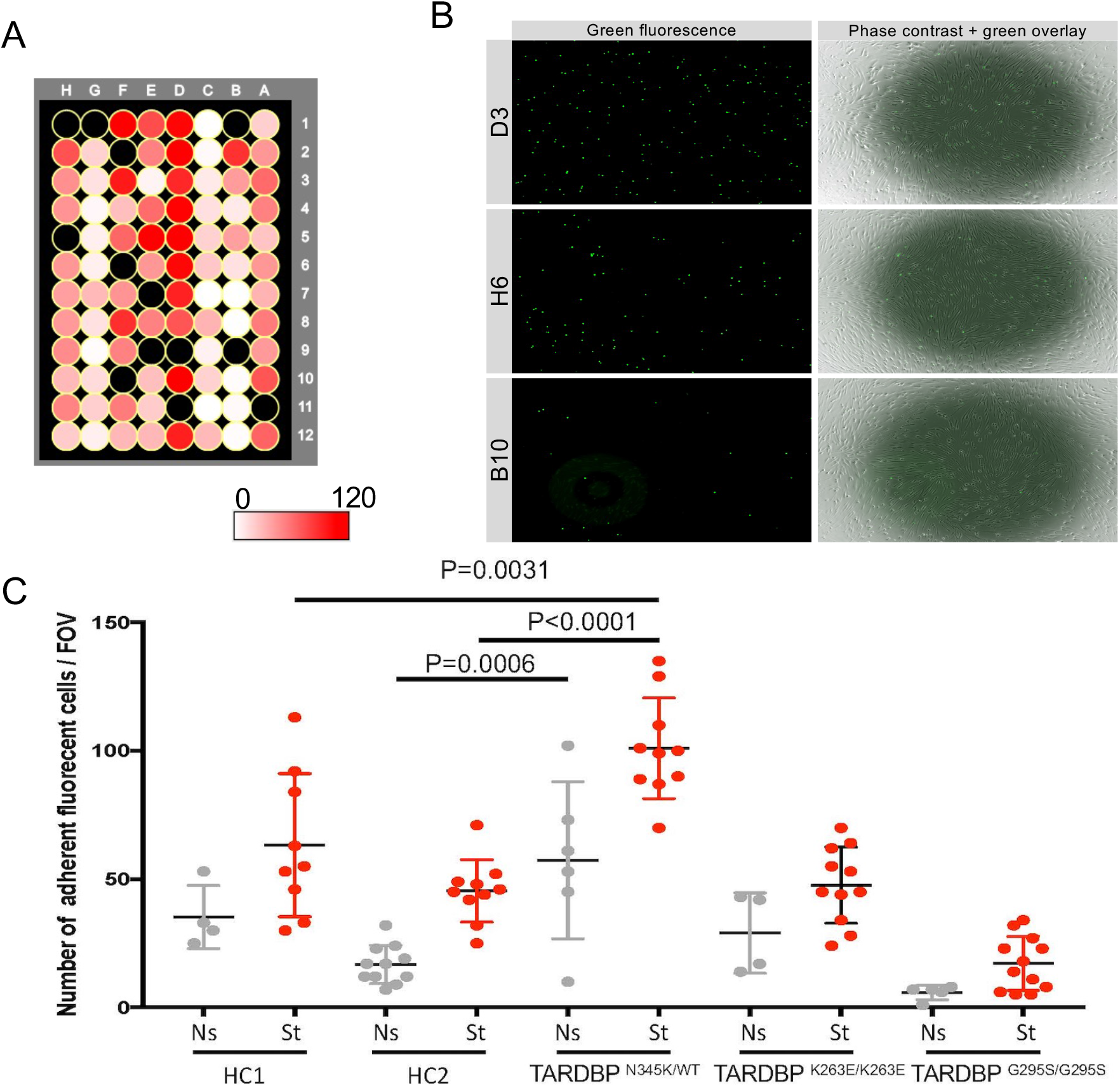
ALS-patient derived EECM-BMEC-like cells enhanced PBMCs adherent. EECM-BMEC-like cells are seeded onto type IV collagen-coated 96-well plates and treated with conditioned medium from the same hiPSC clone, with or without 16 hours proinflammatory cytokines stimulation (0.1 ng/ml TNF-α + 2IU/ml IFN-γ). Fluorescently labeled PBMCs from healthy donors are interacted with the EECM-BMEC-like cell monolayer for 30 minutes. After washing, phase contrast and fluorescence images from the center of each well was photographed automatically. The number of adherent PBMCs is automatically counted using the hybrid cell count software BZ-X analyzer.(A)The heat map of the number of adherent PBMCs in the wells. A1-12 were ALS/FTLD1 under non-stimulated conditions. B1-6 were ALS/FTLD1 under cytokine-stimulated conditions. B7-12 were ALS/FTLD2 under non-stimulated conditions. C1-12 were ALS/FTLD2 under cytokine-stimulated conditions. D1-12 were ALS under cytokine-stimulated conditions. E1-6 were ALS under non-stimulated conditions. E7-12 were HC1 under non-stimulated conditions. F1-12 were HC1 under cytokine-stimulated conditions. G1-12 were HC2 under cytokine-stimulated conditions. H1-12 were HC2 under cytokine-stimulated conditions. Black colored wells were excluded because the cell layers in the center of the wells were detached. (B) Representative images of each well. Green fluorophore-labeled adherent PBMCs and monolayers of EECM-BMEC-like cells on phase contract image. D3, H6, and B10 were the location of the wells representing a high, middle, and low number of adherent PBMCs, respectively. (C) Number of adherent PBMCs onto EECM-BMEC-like cells derived from each clone in the presence or absence of proinflammatory cytokines stimulation. A dot represents the number of adherent PBMCs in the FOV at the center of each well (gray: non-stimulated condition (Ns), red: stimulated condition with 0.1 ng/ml TNF-α and 2 IU/ml IFN-γ(St)). Data are shown as mean ± SD. TStatistical analysis: unpaired *t* test. P-values are indicated in the corresponding figures.

## 4 Discussion

In the current study, we applied our previously established hiPSC-derived BBB model, EECM-BMEC-like cells, which is extremely useful for studying the endothelial cell morphology and the interactions between the BBB and immune cells, to the study of BBB involvement in ALS pathogenesis. This application allowed us to create a novel human ALS patient-derived BBB model. To date, reports of BBB dysfunction in ALS patients have been limited to studies of postmortem autopsy specimens, making it unclear whether BBB dysfunction is the cause of neurodegeneration or merely a secondary consequence. However, several studies of the rodent models have demonstrated the BBB disruption preceding neuronal loss. For example, in G93A human *SOD1* transgenic mice (*SOD1* ^G93A^), which are widely used as animal models for ALS, motor neuron numbers decrease and neurological symptoms appear at 13 weeks of age. However, (1) the leakage of IgG and Prussian blue has already begun at 8 weeks when the number of motor neurons has not yet decreased (Zhong et al., 2008). (2) Increased expression of matrix metalloproteinase-9, a key extracellular matrix degradation enzyme, and decreased staining of tight junction and basement membrane components were observed at 10 weeks (Miyazaki et al., 2011). (3) IgG deposition, increased ICAM-1, and activation of microglia in the spinal cord were observed from the asymptomatic period around day 40-80 (Alexianu et al., 2001). (4) Even at the early stage of the disease (13 weeks), endothelial cells and endfeet of astrocytes are swollen and vascularized in the brainstem and spinal cord (Garbuzova-Davis et al., 2007). Such”pre-symptomatic barrier dysfunction” has also been reported in SOD1 ^G93A^ rats and SOD1 ^G86R^ mice (Nicaise et al., 2009; Milane et al., 2010). BBB leakage and immune cell infiltration into the CNS have also been reported in *TARDBP*-related mouse models, although the studies did not focus on the onset of the alteration (Sasaki et al., 2015; Zamudio et al., 2020). This evidence encouraged us to investigate whether BBB functions are intrinsically altered in human ALS, independent of neurodegeneration, and contribute to disease onset or progression. Indeed, our novel ALS patient-derived BBB model recapitulates two major BBB disruptive mechanisms, 1) increased permeability and 2) enhanced immune cell attraction, and opens an avenue to investigate this challenging hypothesis.

In the present study, an ALS patient-derived EECM-BMEC-like cell (*TARDBP* ^N345K^) exhibited discontinuation of tight junctions resulting in increased permeability of small molecule tracers. This model also showed increased expression of ICAM-1 and VCAM-1, which are major adhesion molecules for promoting immune cell trafficking into the CNS. Moreover, enhanced endothelial adhesion molecules are functional since we showed a higher number of PBMCs adhered to ALS-derived EECM-BMEC-like cells compared to HCs-derived cells. One of the advantages of disease modeling using hiPSCs is that we can simplify the model and investigate how a patient genetic background affects the specific cell type. In this study, we showed that dysfunction of BMEC-like cells occurs independently of the impact of degenerating neurons or other glial cells. Similar to our result, the other group reported on the impaired BBB functions of hiPSC-derived BMEC-like cells from ALS patients harboring *SOD1* and *C9orf72* mutations (Katt et al., 2019), although BMEC-like cells are differentiated using different methods (Lippmann et al., 2014) and cell surface adhesion molecule could not be investigated in this cases (Nishihara et al., 2020). These *SOD1* or *C9orf72* mutation-carrying hiPSC-derived BMEC-like cells showed a lower transendothelial electrical resistance (TEER), but the permeability of the small molecule tracer was not significantly different compared to HCs. It is unclear whether the difference in permeability was due to differences in the mutation or to differences in the method. The conventional model presents a mixed epithelial and endothelial phenotype with a very low permeability (Lippmann et al., 2020), making it difficult to detect differences in permeability. In any case, these data suggest that the BBB in ALS patients may be genetically fragile. The ALS patient harbored heterozygous SNP mutation at the coding exon 6 of *TARDBP*, resulting in a missense mutation (p.N345K) reported at the site known to be involved in the interaction of TDP-43 with other proteins (Rutherford et al., 2008). In this study, we also used *TARDBP* ^G295S/G295S^ and *TARDBP* ^K263E/K263E^ gene-editing hiPSCs, which mutations are known to be pathogenic that cause ALS and/or FTLD (Floris et al., 2015). We have previously reported that differentiated neurons using the same hiPSC lines with *TARDBP* ^K263E/K263E^ show a loss of function of TDP-43, indicating its usefulness in studying the impact of the mutation (Imaizumi et al., 2022). We asked if the specific *TARDBP* mutations contribute to BBB dysfunction, however, *TARDBP*^G295S/G295S^ and *TARDBP*^K263E/K263E^ gene-editing hiPSCs-derived EECM-BMEC-like cells didn’t show impaired diffusion barrier functions nor enhanced immune cell adhesion compared to the isogenic control. The discrepancy between an ALS patient (*TARDBP* ^N345K^) and gene editing isogenic clones (*TARDBP* ^G295S/G295S^ and *TARDBP* ^K263E/K263E^) might be the difference in the site of mutation (Figure 1A) or the type of allele mutation (heterozygous vs. homozygous). Another explanation is that EECM-BMEC-like cells with G295S and K263E mutation might not impair RNA processing as previously reported in nonneuronal cells (Imaizumi et al., 2022).

In the current study, we have also newly established the method of studying immune cell interaction at the BBB in the 96-well plate. The conventional assay could only compare a small number of wells, such as 8 to 16 wells (Nishihara et al., 2020; Nishihara et al., 2021), but the immune cell status could change from experiment to experiment, making it problematic to compare experiments performed on different days. In addition, the distribution of adherent cells is not always equal, and there was a possibility of bias at the imaging site when the analyzer chose where to photograph. To address these issues, we have modified the protocol to create a more effective assay that compares multiple clones under the same conditions in an unbiased manner using an automated analysis system. The major strength of our unique iPSC-derived BMEC-like cells is that they combine robust barrier function with the expression of cell adhesion molecules necessary for immune cell trafficking (Nishihara et al., 2020). Incorporating the iPSC-derived model into this newly established multi-well assay allows us to further accelerate the study of the interaction between the BBB and immune cells using multiple clones.

Our study has limitations. First, we have only analyzed a patient-derived model and cannot conclude whether the specific ALS-associated mutation (*TARDBP* ^N345K^) contributes to BBB dysfunction. Therefore, we cannot exclude the possibility that genetic background is an additional factor in barrier failure independent of *TARDBP* mutations or acts as a co-factor with *TARDBP* mutations necessary to affect BMEC functions. In the future, functional analysis using an isogenic model of *TARDBP* ^N345K^ will be required for confirmation. Second, we have shown in this model an impaired diffusion barrier and the potential of BMECs for immune cell infiltration into the CNS. However, we did not provide evidence that BBB dysfunction directly contributes to neurodegeneration. For future studies, advancing the BBB model, which recapitulates blood flow and incorporates neurons, astrocytes, and microglia is needed to show direct evidence of how peripheral changes cause neurodegeneration.

In conclusion, we established an ALS-derived BBB model that is useful in studying diffusion barrier functions and functional cell surface adhesion molecules. Familial ALS patient-derived BBB model showed disruption of tight junctions resulting in increased permeability of small molecule tracers and increased expression of cell surface ICAM-1 and VCAM-1, functionally attracting immune cells. Using the isogenic model, we show that the specific mutation of TARDBP (*TARDBP* ^G295S/G295S^ and *TARDBP* ^K263E/K263E^) does not contribute to these barrier dysfunctions. Thus, our novel ALS patient-derived BBB model is useful for studying the involvement of BBB dysfunction in the pathogenesis of ALS and promoting therapeutic drug discovery targeting BBB. The novel automated techniques for analyzing immune cell BBB adhesion in multiple samples facilitate further study of immune cell-BBB interactions.

## Author Contributions

K.M., K.N., R.U., T.S., and H. N. experimented and analyzed data. N.S. and M.A. recruited the patient (ALS), S.M. and H.O. provided hiPSC clones (ALS, ALS/FTLD1, and ALS/FTLD2). K.M. and H.N. wrote the manuscript with support from S.M., H.O., and M.N. H.N. conceptualization and supervision of the study, funding acquisition; project administration.

## Acknowledgments

We are grateful to Shiho Nakamura and Fumiko Ozawa (Keio University) for technical assistance, and GenAhead Bio for generating isogenic TDP-43 ^G295S^ and TDP-43 ^K263E^ iPSCs.

