## Supplementary table 2 for "Establishment of a novel amyotrophic lateral sclerosis patient-derived blood-brain barrier model: Investigating barrier dysfunction and immune cell interaction"

| Specific reagent | ingredient | Volume |
|---|---|---|
| Blocking buffer | Skim milk | 5 g |
| | Tris-Bufferwd Saline 1x | 100 mL |
| | Triton X-100 | 300 µL |
| | NaN3, 10% solution in water | 400 µL |
| Human fibroblast growth factor 2 (bFGF/FGF2) stock solution | Human fibroblast growth factor 2 | 500 µg |
| | dPBS | 5 mL |
| | Bovine serum albumin (BSA), 7.5% in dPBS | 66.7 µL |
| hECSR medium | Human Endothelial Serum Free Medium | 500 ml |
| | FGF2 stack solution | 100 µL |
| | B-27 Supplement (503), serum free | 10 ml |
| INFγ stock solution | Recombinant Human IFN-gamma Protein | 100 µg |
| | dPBS | 500 µL |
| | BSA, 7.5% in dPBS | 6.667 µL |

| Specific reagent | ingredient | Volume |
|---|---|---|
| Migration assay medium | DMEM , [+] 4.5 g/L D-Glucose, [-] ʟ-Glutamine, [-] Pyruvate n/a | 90.5 mL |
| | HEPES | 2.5 mL |
| | Fetal Bovine Serum, | 5 mL |
| | L-Glutamine 200mM | 2mL |
| Sodium fluorescein stock solution | dPBS | 2.6577 mL |
| | Fluorescein sodium salt | 10 mg |
| T-cell medium | RPMI Medium 1640 | 82.8 mL |
| | L-Glutamine 200 mM (100x) | 1 mL |
| | Fetal Bovine Serum | 10 mL |
| T-cell wash buffer | T-cell medium | 10 mL |
| | RPMI Medium 1640 | 87.5 mL |
| | HEPES Buffer Solution | 2.5 mL |
| TNFα stock solution | Recombinant Human TNF-alpha Protein | 20 µg |
| | dPBS | 200 µL |
| | BSA, 7.5% in dPBS | 2.667 µL |
| Tris-Buffered Saline (TBS) | Tris Base | 6.05 g |
| | NaCl | 8.76 g |
| | distilled water | ～1000 mL |
