## Supplementary table 3 for "Establishment of a novel amyotrophic lateral sclerosis patient-derived blood-brain barrier model: Investigating barrier dysfunction and immune cell interaction"

| | product name | corporation / catalog number | dilution | fixative |
|---|---|---|---|---|
| For immunostaining | Anti-VE-cadherin antibody (clone F-8), | Santa Cruz Biotechnology / sc-9989 | 1:200 | MeOH for 20 sec |
| | Claudin 5 Monoclonal Antibody (clone 4C3C2) | Invitrogen / 34-1600 | 1:200 | MeOH for 20 sec |
| | Occludin Monoclonal Antibody (clone OC-3F10), | Invitrogen / 33-1500 | 1:100 | MeOH for 20 sec |
| | Actin, Smooth Muscle Ab-1, Mouse Monoclonal Antibody (clone 1A4) | Fisher Scientific / MS113P | 1:200 | 1% PFA |
| | Monoclonal Anti-Calponin antibody produced in mouse (clone hCP) | Sigma-Aldrich/ C2687 | 1:15000 | 1% PFA |
| | Anti-TAGLN/Transgelin (polyclonal) | Abcam / ab14106 | 1:1000 | 1% PFA |
| | Cy3 AffiniPure F(ab') Fragment Goat Anti-Mouse IgG, F(ab')2 fragment | Jackson ImmunoResearch / 115-166-006 | 1:200 | |
| | Cy3 AffiniPure Donkey Anti-Rabbit IgG (H+L), (dilution 1:200) | Jackson ImmunoResearch / 711-165-152 | 1:200 | |
| | DAPI (4',6-Diamidino-2-Phenylindole, Dihydrochloride) | Sigma-Aldrich / D9564 | 1:1000 | |
| For flow cytometry | PE anti-human CD106 Antibody | BioLegend / 305806 | 1:20 | |
| | PE Mouse IgG1, κ Isotype Ctrl | Biolegend / 981804 | 1:160 | |
| | BV421 Mouse Anti-Human CD54 | BD / 564077 | 1:20 | |
| | BV421 Mouse IgG1, k Isotype Control | BD / 562438 | 1:160 | |
